# Landscapes associated with Japanese encephalitis virus reflect the functional biogeography of waterbird species across Australia and the Central Indo-Pacific region

**DOI:** 10.1101/2023.08.18.553798

**Authors:** Michael G. Walsh, Cameron Webb, Victoria Brookes

**Affiliations:** The University of Sydney, Faculty of Medicine and Health, Sydney School of Public Health, Camperdown, New South Wales, Australia; The University of Sydney, Faculty of Medicine and Health, Sydney Infectious Diseases Institute, Westmead, New South Wales, Australia; One Health Centre, The Prasanna School of Public Health, Manipal Academy of Higher Education, Manipal, Karnataka, India; The Prasanna School of Public Health, Manipal Academy of Higher Education, Manipal, Karnataka, India; Department of Medical Entomology, NSW Health Pathology, Westmead Hospital, Westmead, New South Wales, Australia; Sydney School of Veterinary Science, Faculty of Science, The University of Sydney, New South Wales, Camperdown

**Author notes:** Address correspondence to: Michael Walsh, PhD, MPH, Senior Lecturer, Infectious Diseases Epidemiology Sydney School of Public Health, The University of Sydney Edward Ford Building Camperdown NSW 2006.

## Abstract

Japanese encephalitis virus (JEV), a zoonotic, mosquito-borne virus, has broad circulation across the Central Indo-Pacific biogeographical region (CIPBR), which recently expanded dramatically within this region across southeastern Australia over the summer of 2021-2022. Preliminary investigation of the landscape epidemiology of the outbreaks of JEV in Australian piggeries found associations with particular landscape structure as well as ardeid species richness. The ways in which waterbird species from diverse taxonomic pools with substantial functional variation might couple with JEV-associated landscape structure was not explored, and therefore, key questions regarding the landscape epidemiology and infection ecology of JEV remain unanswered. Moreover, given the established presence of JEV within the CIBPR, the extent to which waterbird species pools in JEV-associated landscapes in Australia reflect broader regional patterns in functional biogeography presents a further knowledge gap particularly with respect to potential virus dispersal via maintenance hosts. The current study investigated waterbird species presence, ecological traits, and functional diversity distribution at landscape scale, and how these aligned with confirmed JEV detections in eastern Australia and the wider CIPBR. The results showed that waterbird habitat associated with JEV detection in Australia in 2022 and more widely across the CIPBR over the last 20 years reflects a range of species representing 8 families in 4 orders (ardeids, anatids, rallids, phalacrocoracids, threskiornithids, gruids, and pelecanids). Increasing waterbird functional diversity (trait-based mean pairwise dissimilarity) was associated with landscapes delineating JEV occurrence, while only one individual trait, high hand-wing index, was consistently associated with species presence in these JEV-associated landscapes in both Australia and the broader CIPBR. This suggests that dispersal capacity among the waterbird species pools that dominate JEV-associated landscapes might be important. By taking an agnostic approach to JEV maintenance host status, this study indicates a relatively large, CIPBR-wide pool of waterbird families associated with JEV landscapes, challenging the narrow view that JEV maintenance is limited to ardeid birds. In addition, these findings highlight the potential for leveraging functional biogeography in high-risk landscapes across broad geographic extent to guide landscape-specific selection of species for JEV surveillance.

## Introduction

Japanese encephalitis virus (JEV) has circulated in the Central Indo-Pacific biogeographical region (CIPBR) for many decades(Garjito et al., 2018; Hanson et al., 2004; Johansen et al., 2000; Lopez et al., 2015). In early 2022 Japanese encephalitis virus (JEV) was detected for the first time in south-eastern regions of Australia, resulting in the declaration of a Communicable Disease Incident of National Significance. A total of 45 cases of human disease were reported, including seven fatalities(Chief Medical Officer, 2023). Outbreaks were also reported in piggeries with unprecedented wide geographic distribution, high numbers of affected piggeries, and high incidence within piggeries (Australian Department of Health and Aged Care, 2022; Mackenzie et al., 2022; Williams et al., 2022). Previous JEV occurrence in Australia had been limited to isolated cases in the Torres Strait Islands(Hanna et al., 1996) and the Cape York Peninsula of northern Queensland(Hanna et al., 1999). The extensive geographic distribution of the recent outbreaks, which included eastern and southern Australia, was of particular concern because it deviated markedly from anticipated emergence events for JEV(van den Hurk et al., 2019) and suggested broad circulation of the virus in maintenance communities prior to emergence in pigs and humans(Purnell, 2022). These outbreaks followed a long period of anomalous weather due to consecutive La Niña phases of the El Niño Southern Oscillation (ENSO), including high levels of precipitation and major flooding events across eastern Australia that generated considerable disturbance to natural communities(Bureau of Meteorology, 2021). It was hypothesised that the anomalous La Niña-associated weather patterns might have contributed to the widespread distribution of JEV outbreaks in 2022 due to modulation of vector or maintenance host population distributions(Purnell, 2022). However, a thorough investigation of landscape composition and configuration and La Niña-associated anomalies showed that landscape structure was substantially more influential to outbreak occurrence than were specific precipitation or temperature anomalies associated with La Niña(Walsh et al., 2023). Aspects of high-risk landscape structure, such as the composition of transient wetlands and the propensity to flooding, would also have been affected by both proximal and distal precipitation events, and so La Niña-associated weather anomalies likely influenced outbreak occurrence indirectly.

Japanese encephalitis virus circulates in wild ardeid bird maintenance hosts(Bhattacharya and Basu, 2014; Buescher et al., 1959; Jamgaonkar et al., 2003; Nemeth et al., 2012; Ogata et al., 1970; Rodrigues et al., 1981; Scherer, 1959) vectored by mosquitoes (primarily *Culex annulirostris* in Australia(van den Hurk et al., 2003) and *Cx. tritaeniorhynchus* in Southeast Asia(Endy and Nisalak, 2002; Longbottom et al., 2017)) and can spillover to pigs(Baruah et al., 2018; Borah et al., 2013; Chen and Beaty, 1982; Ghimire et al., 2014; Kakkar et al., 2017; Komada et al., 1968) and humans, which are competent and non-competent hosts, respectively. Previous work has examined the distribution of ardeid species richness as a key nidus of the maintenance community, identifying a strong quadratic relationship between richness and JEV piggery outbreak occurrence, in which intermediate richness was associated with greatest risk(Walsh et al., 2023). However, only ardeid species richness was explored, and not the association between outbreaks and individual species from a broader taxonomic pool, or the influence of the structure and functional distribution of the species pool on the 2022 piggery outbreaks in Australia. The latter has not been assessed with respect to outbreaks anywhere in the CIPBR to date, despite evidence of JEV circulation across the Torres Strait Islands(Johansen et al., 2004; van den Hurk et al., 2001) and far north Queensland(van den Hurk et al., 2006), as well as in Papua New Guinea(Hanson et al., 2004; Johansen et al., 2000), for 2 decades, and in Indonesia, Malaysia, and the Philippines for several decades(Garjito et al., 2018; Lopez et al., 2015). Due to the logistics and resources required to sample birds across the vast landscapes of both Australia and the broader CIPBR, as well as the potential for extensive dispersal of maintenance hosts and vectors within the CIPBR, a better understanding of how particular species and species pools align with landscapes associated with JEV occurrence would be useful to identify target species to monitor JEV circulation in these landscapes. This could be especially useful to illuminate the full breadth of potential JEV maintenance hosts, given that a narrower perspective has typically limited the scope of wild host sampling to ardeids and recent studies have provided evidence that this scope should be expanded(Levesque et al., 2024; Moore et al., 2024).

The study of the functional ecology of viral maintenance hosts has increased in recent years as a means to help identify potential hosts based on specific biological and ecological traits, frequently focusing on life history due to the association between fast-living and host competence in many r-selected species(Brunner et al., 2008; Cynthia J. Downs et al., 2019; Cynthia J Downs et al., 2019; Han et al., 2016, 2015a, 2015b; Ostfeld et al., 2014; Plourde et al., 2017; Stephens et al., 2016). These species, which are typically generalist and frequently more resilient to anthropogenic pressure(Faust et al., 2018; Gibb et al., 2020; Johnson et al., 2020), tend to invest more in earlier and more fecund reproductive effort, and less in immune function and adult survival(Molles, 2015). As such, many of these species are also optimal hosts for pathogens(Johnson et al., 2012). Nevertheless, the association between fast-living species and host competence is not universal, with some systems demonstrating the opposite or ambiguous associations, such as ebolaviruses(Schmidt et al., 2019), Rift Valley fever virus (RVFV)(Walsh and Mor, 2018), and Ross River virus (RRV)(Walsh, 2019). In addition to life history, other host traits, including population density, diet composition, and dispersal ability, can also be influential to the infection ecology of some pathogens(Gibb et al., 2020; Johnson et al., 2020). Critically, relationships with species traits, particularly those relating to life history and population-level transmission dynamics, have been largely neglected in bird-host pathogen systems. However, a recent example demonstrated strong associations between wild avian influenza hosts and life history and dispersal capacity, which is another widely distributed, albeit not vector-borne, host-virus system closely associated with agricultural-wetland mosaics(Yin et al., 2023). Finally, functional diversity offers a further key representation of ecological diversity, which can help to inform ecological relationships beyond what is demonstrated by community composition (i.e. species richness, evenness, abundance) or phylogenetic diversity alone. The extent to which landscape-scale species pools diverge or converge in their biological and ecological traits can help to inform environmental filtering, dispersal capacity, and biotic interactions, particularly in the context of JEV occurrence. Notwithstanding the importance of species traits to species distributions, host functional ecology has hitherto gone unexplored with respect to JEV hosts and is, therefore, a critical knowledge gap for this system especially since the functional biogeography of waterbirds in Australia and the broader CIPBR might be directly relevant to JEV circulation across the region.

The first aim of this study was to identify how species pools mapped to JEV-associated landscapes in Australia. Specifically, we sought to determine which waterbird species (eight families of 4 avian orders) were distributed in close association with JEV piggery outbreaks, and how species pools were structured with the landscape composition and configuration of these outbreaks, to inform preliminary target host species in specific landscapes for One Health surveillance of JEV. Secondly, we aimed to determine the extent to which species pool-weighted trait means and functional diversity were coupled with JEV-associated landscapes in Australia. Finally, we aimed to determine whether species pools and their functional ecology similarly mapped to landscapes where JEV has been detected across the broader CIPBR as a whole.

## Material and methods

### Data Sources

#### Animal data

Australian piggery outbreaks (N = 54) reported to the World Organisation for Animal Health (WOAH) by the Australian government between 19 January, 2022 and 4 April, 2022(World Organisation For Animal Health, 2022) have been described in a previous epidemiological investigation(Walsh et al., 2023). Briefly, outbreak occurrence was defined as one or more cases reported at the level of the piggery. An additional 11 independently-documented JEV occurrence locations (eight piggery outbreaks reported by a commercial pork producer, and 3 positive mosquito pools detected through the NSW Arbovirus Surveillance and Mosquito Monitoring Program(New South Wales Health, 2022)) were used to externally validate model performance. For the extended CIPBR analyses, 25 additional JEV occurrence detections (9 mosquito pool detections, 14 human detections, and 2 pig detections) in Papua New Guinea (n = 4)(Hanson et al., 2004; Johansen et al., 2000), the Torres Strait Islands (n = 2)(Johansen et al., 2004; van den Hurk et al., 2001), the Cape York Peninsula in Australia (n = 1)(van den Hurk et al., 2006), Indonesia (n = 4)(Faizah et al., 2021; Kardena et al., 2024; Spicer et al., 1999), Malaysia (n = 12)(Khor et al., 2020), and the Philippines (n = 2)(Aure et al., 2022) were identified from laboratory confirmed investigations that were published between 1997 and 2019. Geographic locations were published along with these data and latitude and longitude coordinates were obtained from Google Maps when not otherwise specifically provided. All reporting of these detections was available at a minimum of 2.5 arc minutes resolution.

The habitat suitability of extant species of waterbirds across the CIPBR was modelled using 5,190,722 observations of these species representing the Ardeidae, Threskiornithidae, Pelecanidae, Anatidae, Rallidae, Phalacrocoracidae, Ciconiidae, Gruidae avian families that are either extant to mainland Australia (n = 69 species), or the broader the CIPBR without being extant to Australia (n = 23 species), for a total of 92 species currently described throughout the region. Observations were recorded between 1 January 2010 and 31 December 2020 and acquired from the Global Biodiversity Information Facility (GBIF)(Global Biodiversity Information Facility, n.d., n.d.). The phylogenetic tree of these species was obtained from the VertLife project using the Hackett backbone and constructed based on the methods described by Jetz et al. (Jetz et al., 2012). Taxonomic binomial discrepancies between the VertLife tree and the named species obtained from GBIF used for the species distribution models described below were resolved and alternate binomials appear in parentheses in Table S1, with one exception. *Porphyrio melanotus* and *Porphyrio indicus* were previously considered subspecies of *Porphyrio porphyrio*, and so the former 2 species are represented by the latter binomial, *Porphyrio porphyrio*, in the phylogenetic tree. Thus, all phylogenetic analyses comprise 91 species rather than the current full 92 species. Data obtained from the national pig herd dataset, which is used in the Australian Animal Disease Spread Model, were used to quantify Australian domestic pig density for the JEV models described below. This dataset comprises >8000 registered pig herds of all types (including commercial, boar studs, smallholder and pig keepers)(Bradhurst et al., 2015). For the rest of the CIPBR, pig density was obtained from the Gridded Livestock of the World database(Robinson et al., 2014).

Reporting bias may affect JEV occurrence detection, whereby large commercial herds may be more likely to capture and record cases due to more systematic collection of production data and increased probability of disease detection in larger herds. Moreover, all JEV detections in this study were associated with piggeries either directly or indirectly, from the piggery outbreak data in Australia, to human outbreaks proximate to piggeries and pig sentinel and mosquito vector surveillance in other parts of the CIPBR. Therefore, background points for JEV models were selected proportional to pig density throughout the CIPBR to correct for potential JEV detection reporting bias.

#### Environmental data for habitat suitability models

Land cover classification based on the European Space Agency and Climate Change Initiative was used to classify wetlands(European Space Agency and Climate Change Initiative, 2017a; Lamarche et al., 2017) and land cover(European Space Agency and Climate Change Initiative, 2017b) at 3 arc seconds in 2010, as this time point corresponded to the beginning of the period of recorded ardeid observations described above. Waterways were represented by a separate high-resolution (3 arc seconds) image produced in collaboration between CIESIN and the WorldPop project (University of Southampton et al., n.d.; WorldPop, n.d.). Mean annual precipitation, mean annual temperature, and isothermality were obtained from WorldClim (Fick and Hijmans, 2017). The Priestley-Taylor α coefficient (P-Tα) is the ratio of actual evapotranspiration to potential evapotranspiration and was used to describe water stress in the landscape(Khaldi et al., 2014; Priestley and Taylor, 1972). The P- Tα data were acquired from the Consultative Group for International Agricultural Research (CGIAR) Consortium for Spatial Information at a resolution of 30 arc seconds(Trabucco and Zomer, 2010).

#### Environmental data for landscape-level metrics (Australia-only)

All JEV outbreaks in Australia were reported to a scale of at least 5 arc minutes (∼ 10 km), which thus represents the landscape-level scale of the current investigation. Land cover classes were represented using the previously validated Copernicus land cover data at 100 m resolution(Jung et al., 2020a, 2020b). The Joint Research Centre’s Global Surface Water product (based on Landsat 5, 7, and 8 imagery) was used to represent high resolution (30 m) surface water seasonality in the year prior to piggery outbreaks(Joint Research Centre, n.d.; Pekel et al., 2016). Each pixel in the raster tiles represents the total number of months (0 – 12) with surface water present during 2021. The heterogeneity of water transience across the landscape was based on five derived rasters: 1) permanent surface water presence over the 12 months of 2021, 2) surface water absence over the 12 months of 2021, 3) surface water presence for 1 to 3 months of the year, 4) surface water presence for 4 to 6 months of the year, and 5) surface water presence for 7 to 9 months of the year. Water movement through and accumulation in the landscape was quantified using hydrological flow accumulation, which describes the total upland area draining into each 500m by 500m parcel and was acquired from the Hydrological Data and Maps based on SHuttle Elevation Derivatives at multiple Scales (HydroSHEDS) information system(Lehner et al., 2006).

#### Waterbird trait data

Bird species’ diet composition and body mass were obtained from the Elton Traits database(Wilman et al., 2014). Diet composition may be an important characteristic of host species because this can delineate generalist versus specialist species and potential resilience to human pressure(Swihart et al., 2003). Only the dietary proportions of fish and invertebrates demonstrated heterogeneity among the ardeid species, so only these dietary elements were investigated. Body mass is an important life history characteristic as well as being a potential modulator of immune function(Cynthia J Downs et al., 2019; Cynthia J. Downs et al., 2019). Body mass was log transformed prior to analysis. The TetraDENSITY database was used to obtain species mean population density(Santini et al., 2018), the AVONET database was used to obtain the hand-wing index(Tobias et al., 2022), and the Lislevand bird traits database was used to obtain the reproductive characteristic, egg mass(Lislevand et al., 2007). These species traits were included since population density may influence the dynamics of JEV circulation in populations and communities(Adler et al., 2008; Begon et al., 2002; Rodríguez-Pastor et al., 2019; Thrall et al., 1995), hand-wing index has been shown to be the best metric to represent bird dispersal(Kennedy et al., 2016; Lockwood et al., 1998) particularly with respect to dispersal in response to climate and habitat loss(Sheard et al., 2020; Weeks et al., 2023) and has been associated with other avian viral outbreaks previously(Yin et al., 2023), and egg mass is an important indicator of reproductive life history. The data for population density, hand-wing index, and egg mass were missing for 78%, 23%, and 29% of species, respectively, so a random forest machine learning algorithm, which was previously shown to be a robust approach with low error to this application(Fountain-Jones et al., 2019; Penone et al., 2014), was used to impute the missing data. The missForest package was used to implement the algorithm(Stekhoven, 2022).

### Statistical Analysis

#### Species distribution modelling

The habitat suitability of 92 waterbird species present in the CIPBR (69 extant in mainland Australia, 23 extant in the CIPBR but not in Australia) was modelled using the GBIF data described above (Table S1). Habitat suitability models were based on ensembles of boosted regression trees (BRT), random forests (RF), and generalised additive models (GAM), each with spatial block cross- validation at fine-scale (30 arc seconds). The calibration areas for the models were based on a buffer of 500 km for the species presence points. Geographically structured spatial partitions were used to select background points. Observation data were thinned so that only one observation per pixel was included in the analysis to avoid overfitting (Table S1). Environmental features in each suitability model comprised mean annual temperature, mean annual precipitation, isothermality, P-Tα, and proximity to surface water, forest, shrubland, grassland, aquatic vegetation, and cropland. Each model was evaluated according to the model’s omission rate, true skill statistic, and the area under the receiver operating characteristic curve (AUC). Suitability estimates for each species were then derived from an ensemble of the three models (BRT, RF, and GAM) using their average maximum sensitivity and specificity.

Subsequently, the species pool available for community assembly was estimated with each species as follows. Each 1 km^2^ pixel of each species’ habitat suitability estimate that was greater than or equal to the true skill statistic (TSS)(Allouche et al., 2006) was classified as present and summed across all pixels within the 5 arc minute landscape parcel. The TSS was used to designate each parcel as present or absent based on the habitat suitability because it has been shown to be independent of prevalence(Allouche et al., 2006). This summed metric thus provided an estimate for landscape-level species presence available to community assembly for each species from within the species pool at each particular landscape parcel. For each species, this “species pool presence” (SPP) could range from 0, indicating the species was not expected to be present in any 1 km^2^ pixels within the 5 arc minute landscape parcel, to 100 indicating the species was expected to be present in all 1 km^2^ pixels within the landscape parcel. The SPP was thus calculated for each of the 92 waterbird species interrogated and represents the metric used to denote species presence at landscape scale, and was used for “abundance” weighting of the mean pairwise dissimilarity described below although this is not intended to represent species abundance. Direct measurement of community composition, including species abundance and relative abundance, was not possible in the current study since community structure was not observed directly at the appropriate local scale. Instead, the SPP provides the degree of available habitat for each species from within the pool at each landscape parcel. The flexsdm package provided a complete streamlined framework to construct the calibration areas and spatial block partitions, select background points, fit all species distribution models, and derive and evaluate the habitat suitability ensembles as described above(Velazco et al., 2022).

#### Landscape metric analysis (Australia-only)

The metrics used to delineate high-risk landscape structure in the Australian context have been described in detail previously in an analysis of the landscape composition and configuration associated with Australian JEV outbreaks in piggeries(Walsh et al., 2023). Briefly, increasing cultivated land and fragmented grassland were both associated with JEV outbreaks and these were computed as follows. Total cultivated land composition was computed by summing the area of all crop patches in each landscape parcel (landscape-level scale of 5 arc minutes). Grassland fragmentation was assessed by measuring mean patch edge relative to patch area for all grassland patches in each parcel using the related circumscribing circle (RCC). The RCC metric has been shown to be robust to patch size(Csorba and Szabo, 2012) in contrast to the perimeter-to-area ratio (PAR), wherein similarly shaped patches can give different values for different sizes and can translate to significant error when summarising PAR at landscape scale(Csorba and Szabo, 2012). These composition and configuration metrics were previously computed for all land cover classes as referenced above, but only those shown to be associated with JEV outbreak occurrence are investigated with respect to ardeid species here as this was the stated aim of the current study. The landscapemetrics package(Hesselbarth et al., 2019) was used to calculate RCC in R v. 4.3.1.

#### Functional metrics for species traits

As a crude measure of the species pool-weighted trait means at landscape scale, the commonly used community-weighted mean, was calculated for each trait among all species identified as present (as described above under habitat suitability modelling) within each landscape parcel(Laliberte and Legendre, 2010; Lavorel et al., 2008). These “pool-weighted means (PWMs)” are not intended to represent local community trait means, but rather to identify whether any species traits shared among the pool at landscape scale stood out as potentially influential for community assembly (though assembly itself is not described), and particularly in high-risk JEV landscapes. Mean pairwise dissimilarity (MPD) was also computed as a robust metric of functional diversity to evaluate the species pool trait heterogeneity, again quantified at landscape scale and extending across the CIPBR. The MPD was “abundance”-weighted, though, as described above, this weighting was not based on actual estimates of community-level species abundances but rather based on the number of 1 km pixels in which each species was designated present within the 10 km landscape parcel. The FD package(Laliberté et al., 2014) was used to compute weighted means for species traits, the picante package(Kembel et al., 2010) was used to compute MPD, and phytools was used to compute the phylogenetic signal (both Blomberg’s K and Pagel’s λ) of species traits and map continuous traits onto the phylogenetic tree(Revell, 2012) in R v. 4.3.1.

#### Point process modelling

Each of the 92 individual waterbird species’ landscape-level presence within the pool was interrogated with respect to JEV detection by fitting inhomogeneous Poisson point process models (PPMs)(Baddeley et al., 2015; Baddeley and Turner, 2000). To control for potential reporting bias in outbreak surveillance, the background points used in these models were sampled proportional to mean pig density, as described above. For the individual waterbird species PPMs, a simple univariate PPM of JEV outbreaks was created for each of the 92 species. Similarly, for the species trait PPMs, a simple univariate PPM of JEV detections was created for each species trait’s PWM; however, this univariate trait analysis was followed by a multiple PPM to determine which traits among the species pool were independently associated with JEV outbreaks. For the CIPBR models, correlation between the spatial distribution of traits was low (Pearson’s r ≤ 0.66) and all variance inflation factors were ≤ 4.65, indicating collinearity was not a concern for the trait multiple PPM. However, for the Australia-specific model the variance inflation factors indicated multicollinearity with hand-wing index and underwater foraging strategy, so these two variables were fit under separate model structures. All PPMs were fit at a scale of 5 arc minutes, which was the optimum scale of JEV reporting for all detections included in Australia and across the CIPBR(Walsh et al., 2023). The Akaike information criterion (AIC) was used to evaluate model fit. The spatstat R package was used to fit the PPMs(Baddeley and Turner, 2005).

#### Hierarchical modelling of species pools (Australia only)

A hierarchical modelling of species communities (HMSC) framework(Ovaskainen and Abrego, 2020; Tikhonov et al., 2020) was used to assess species pools in relation to the specific features of landscape composition and configuration that have been shown to delineate JEV outbreak risk in the Australian context (Figure S1)(Walsh et al., 2023). In addition, the HMSC framework accounted for the potential modulation of species presence in high-risk landscape structure by their phylogeny and functional traits, associations with the latter representing species “responses” to the landscape features. However these associations do not represent estimates of actual responses within communities and so this interpretation must be tempered (see below). Models were fit under two landscape domains: the first comprised each of the individual landscape features shown to be associated with JEV outbreaks in Australian piggeries (Figure S1) and the second comprised Australian JEV outbreak occurrence itself as the single landscape feature of interest. The two separate domains were evaluated since the landscape features associated with JEV outbreaks did not explain all of the variability in JEV outbreak occurrence, indicating that there are additional unmeasured landscape features that cannot be captured by the first domain, but which may still be relevant to species presence in the landscape. The models were fit with the probit link function to represent species presence, rather than species abundance, as the former is more appropriate given the focus of investigation at landscape scale as already described above. The probit function is preferred over the logit function in HMSC models because it provides a more pragmatic fit to the hierarchical multivariate data(Ovaskainen and Abrego, 2020). These models are fitted using Markov chain Monte Carlo methods and were run here with four chains and 1000 samples each and a burn-in set to 20% of the initial samples. The adjusted r^2^ was used to assess model fit for each of the two model domains. Model convergence was demonstrated for each species (Gelman diagnostic ≈ 1 for each species). Although interspecific interaction within communities was not observed in this study, the HMSC framework does allow for further accounting of important phylogenetic and functional influence on the sharing of space between sympatric species given environmental filtering, which cannot be evaluated with the PPMs described above. Nevertheless, it must be noted that since biotic filtering has not been directly observed, which necessarily occurs at the level of the local community, true community-level functional responses to environmental conditions are not captured at this landscape scale. Instead, these multivariate analyses allow correlative associations to be identified that are anticipated to be useful in characterising how community assembly could potentially operate in landscapes associated with JEV outbreaks, and thereby help to inform sampling targets within such landscapes so that community monitoring of JEV hosts can be systematically structured for One Health surveillance across the large geographic extent of eastern Australia. The Hmsc package(Tikhonov et al., 2020) was used for the HMSC models.

## Results

The PPMs identified landscape-level distributions of 42 species that were associated with Australian JEV outbreaks (7 ardeids, 16 anatids, 8 rallids, 5 phalacrocoracids, 5 threskiornithids, 1 gruid, and 1 pelecanid)(Table S2). When considering the CIPBR as a whole with additional JEV detections in the Philippines, Malaysia, Indonesia, Papua New Guinea, and the Cape York Peninsula, 49 species of the total 92 waterbirds species evaluated were associated with JEV-associated landscapes, again representing each of the eight avian families (Table S3). Of all the individual species traits interrogated under the PPMs, only the PWM distributions of hand-wing index (RR = 1.273; 95% C.I. 1.134 – 1.428) and underwater foraging (RR = 1.05; 95% C.I. 1.014 – 1.088) demonstrated independent associations with the distribution of JEV outbreaks in the Australian context (Table S4, Table 1 and Figure 1), whereas hand-wing index alone (RR = 1.17; 95% C.I. 1.100 – 1.246) was independently associated with increasing JEV detections across the CIPBR as a whole (Table S4, Table 1 and Figure 2). In addition, mean pairwise dissimilarity was positively associated with JEV detections in both Australia and the broader CIPBR, indicating that more functionally diverse species pools were associated with JEV detections. Target maps of high suitability for Australian waterbird sampling for JEV One Health surveillance based on the two PPMs (one incorporating MPD and one incorporating foraging strategy) is presented in Figure 3.

**Figure 1.**
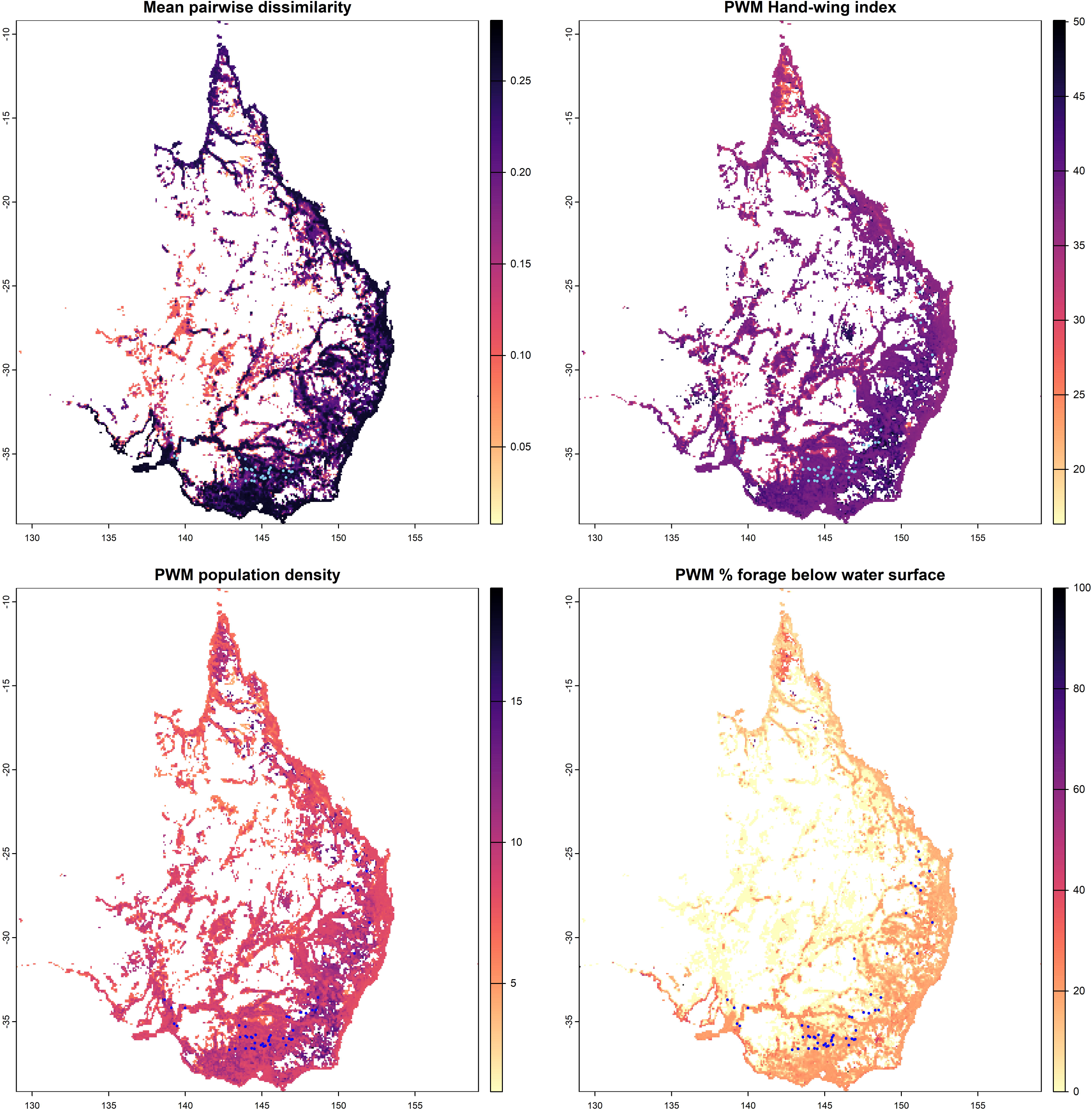
The distribution of the pool-weighted means (PWMs) of species traits for those traits that were associated with Japanese encephalitis outbreaks in the univariate point process models (proportion underwater foraging strategy, hand-wing index, and population density) and mean pairwise dissimilarity (MPD) across Australia. The overlaid points represent the locations of the JEV piggery outbreaks.

**Figure 2.**
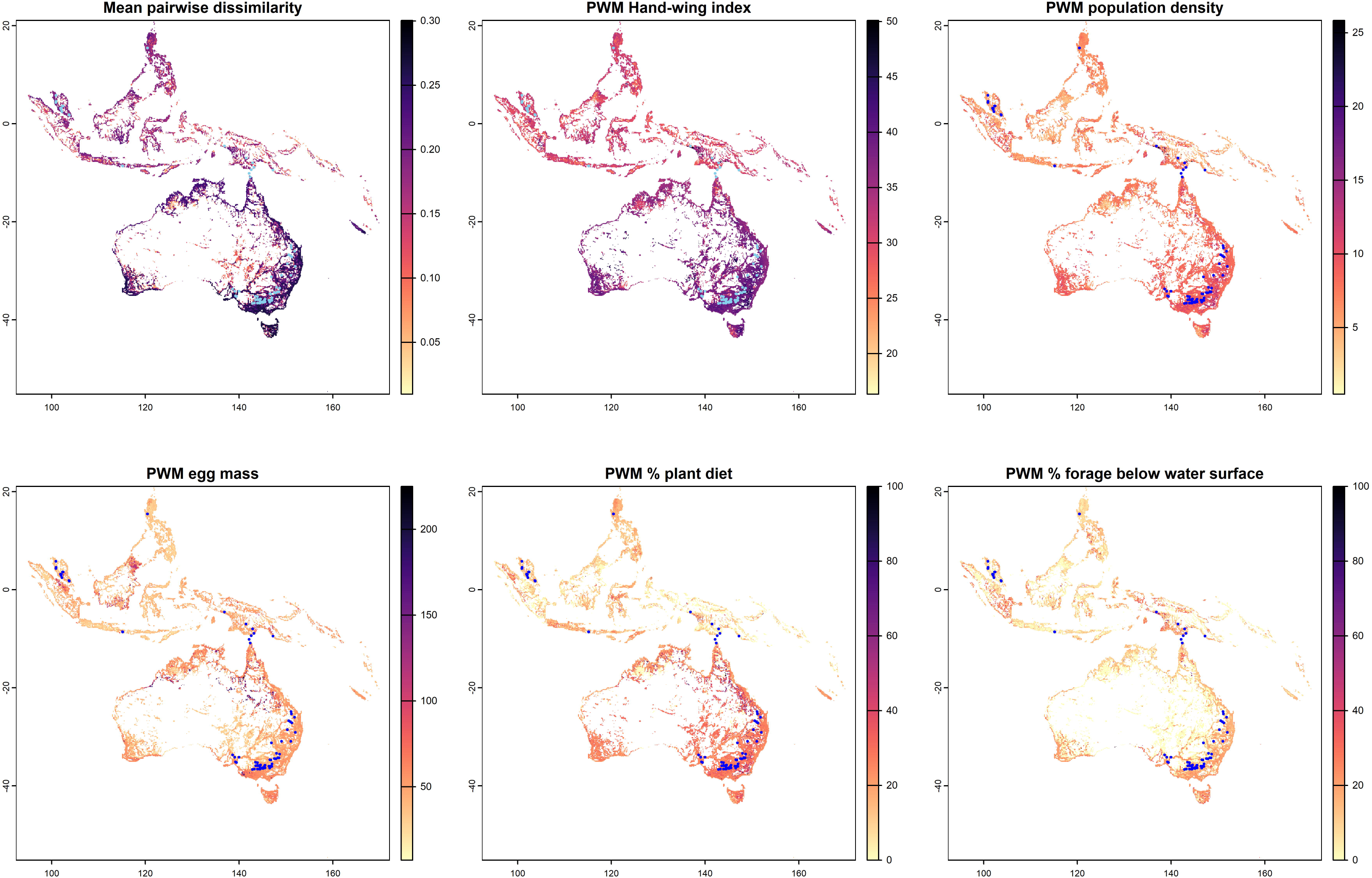
The distribution of the pool-weighted means (PWMs) of species traits for those traits that were associated with Japanese encephalitis outbreaks in the univariate point process models (egg mass, proportion of the diet comprising plants, proportion underwater foraging strategy, hand-wing index, and population density) and mean pairwise dissimilarity (MPD) across the Central Indo-Pacific biogeographical region. The overlaid points represent the locations of the JEV piggery outbreaks.

**Figure 3.**
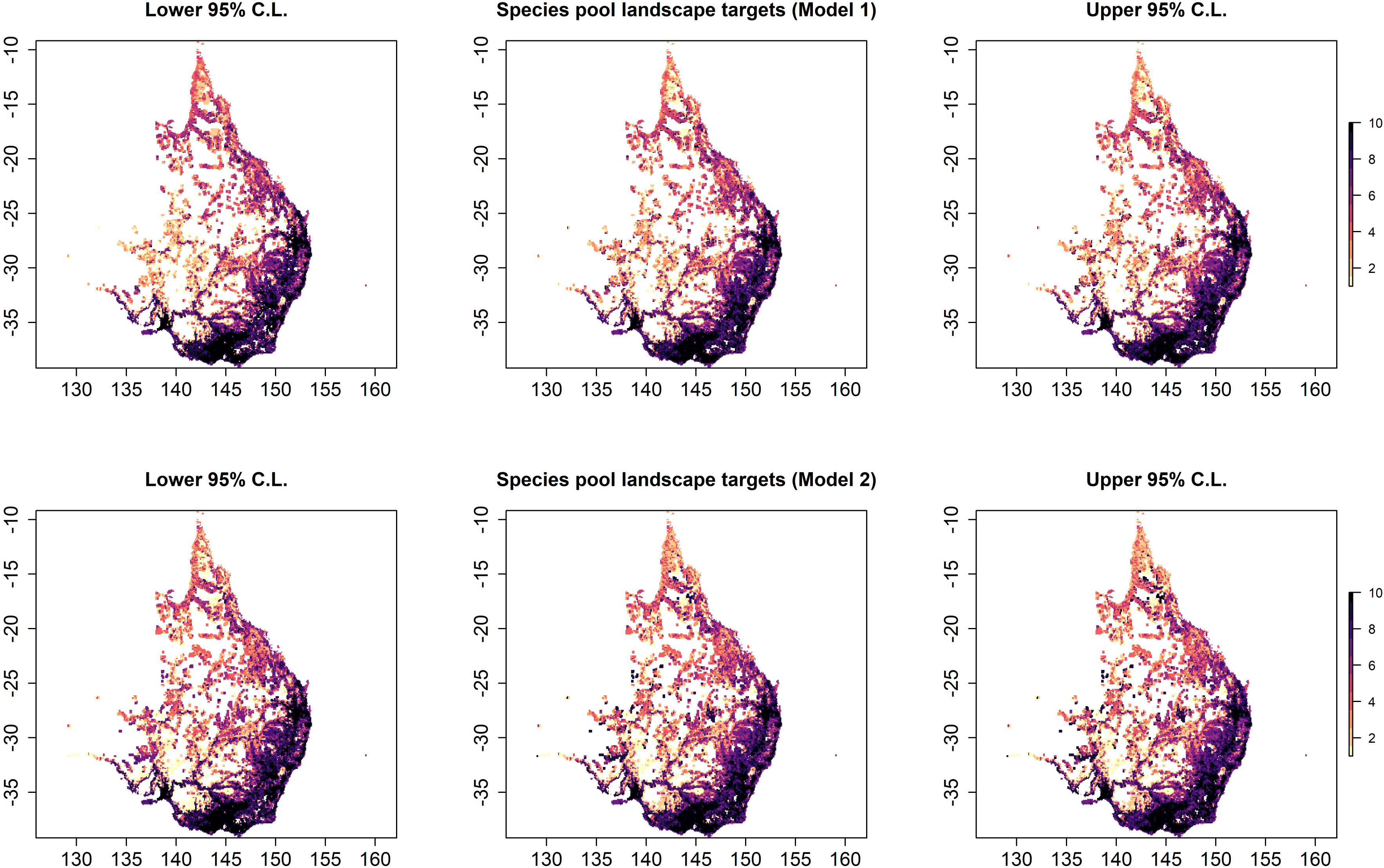
Suggested landscape sampling for Australia based on JEV outbreak intensity estimates as a function of waterbird species traits. The distribution of JEV predicted intensity is presented in the centre panels by decile, whereas the left and right panels represent the lower and upper 95% confidence limits for the predicted intensities, respectively. Predictions are based on the best fitting and performing inhomogeneous Poisson point process model (Table 1A).

**Table 1.** Regression coefficients and 95% confidence intervals for the associations between Japanese encephalitis virus outbreaks and species traits as derived from multiple inhomogeneous Poisson models for Australia alone (A) and the Central Indo-Pacific biogeographical region (CIPBR) as a whole (B).

| Landscape species pool weighted traits | AIC | Estimate | 95% confidence interval |
| --- | --- | --- | --- |
| **A. Australia-specific** |  |  |  |
| *Null model* | 236.75 |  |  |
| *Model 1* | 212.06 |  |  |
| Hand-wing index |  | 1.297 | 1.158 – 1.453 |
| Mean pairwise dissimilarity |  | 9.312 | 2.205 – 16.419 |
| *Model 2* | 213.91 |  |  |
| Hand-wing index |  | 1.273 | 1.134 – 1.428 |
| Forage strategy – underwater |  | 1.050 | 1.014 – 1.088 |
| **B. CIPBR** |  |  |  |
| *Null model* | 838.56 |  |  |
| Hand-wing index | 479.47 | 1.17 | 1.100 – 1.246 |
| Mean pairwise dissimilarity |  | 2.302 | 1.127 – 4.704 |

The Australian-specific HMSC model was in close agreement with respect to the structure of the species pool, identifying similar species presences associated with JEV occurrence as those identified by the PPMs(Figure 4). The median adjusted r^2^ for the landscape feature-based model and JEV-based model was 15.8% and 25.0%, respectively. The HMSC model showed that whilst some high-risk landscape features were common to almost all waterbird species (the presence of transient wetlands and proximity to waterways, as expected), other high-risk features reflected much more heterogenous species pools (Figure 4). By and large, species presence within these high-risk landscape features did not appear closely associated with their functional traits after having accounted for phylogenetic structure, with one exception. There was a distinct association between species presence in landscapes of high water flow accumulation and higher hand-wing index (left panel, Figure 5). Furthermore, interrogating landscapes defined solely by their JEV outbreak occurrence, rather than the individual landscape features associated with outbreaks, showed that species presence in landscapes characterised by outbreak occurrence was also associated with greater species hand-wing index (right panel, Figure 5).

**Figure 4.**
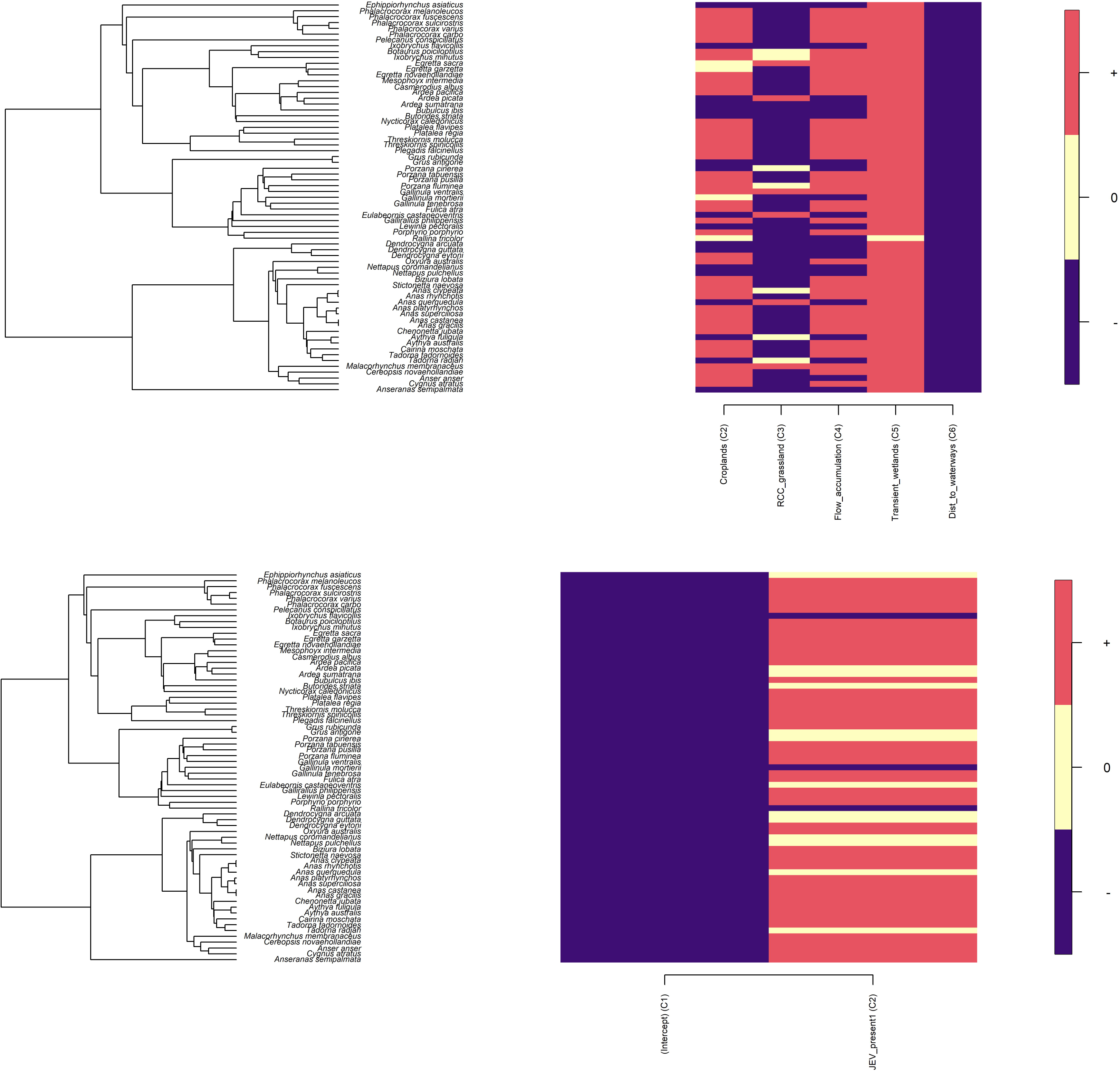
The associations between each species and the landscape features under two model structures using hierarchical modelling of species communities (HMSC). The first HMSC model (top panel) shows associations as positive, negative, or neutral “responses” of each species on the y-axis to each landscape feature associated with JEV outbreaks on the x-axis. The second HMSC model (bottom panel) shows these associations for JEV outbreak occurrence only and no other landscape features.

**Figure 5.**
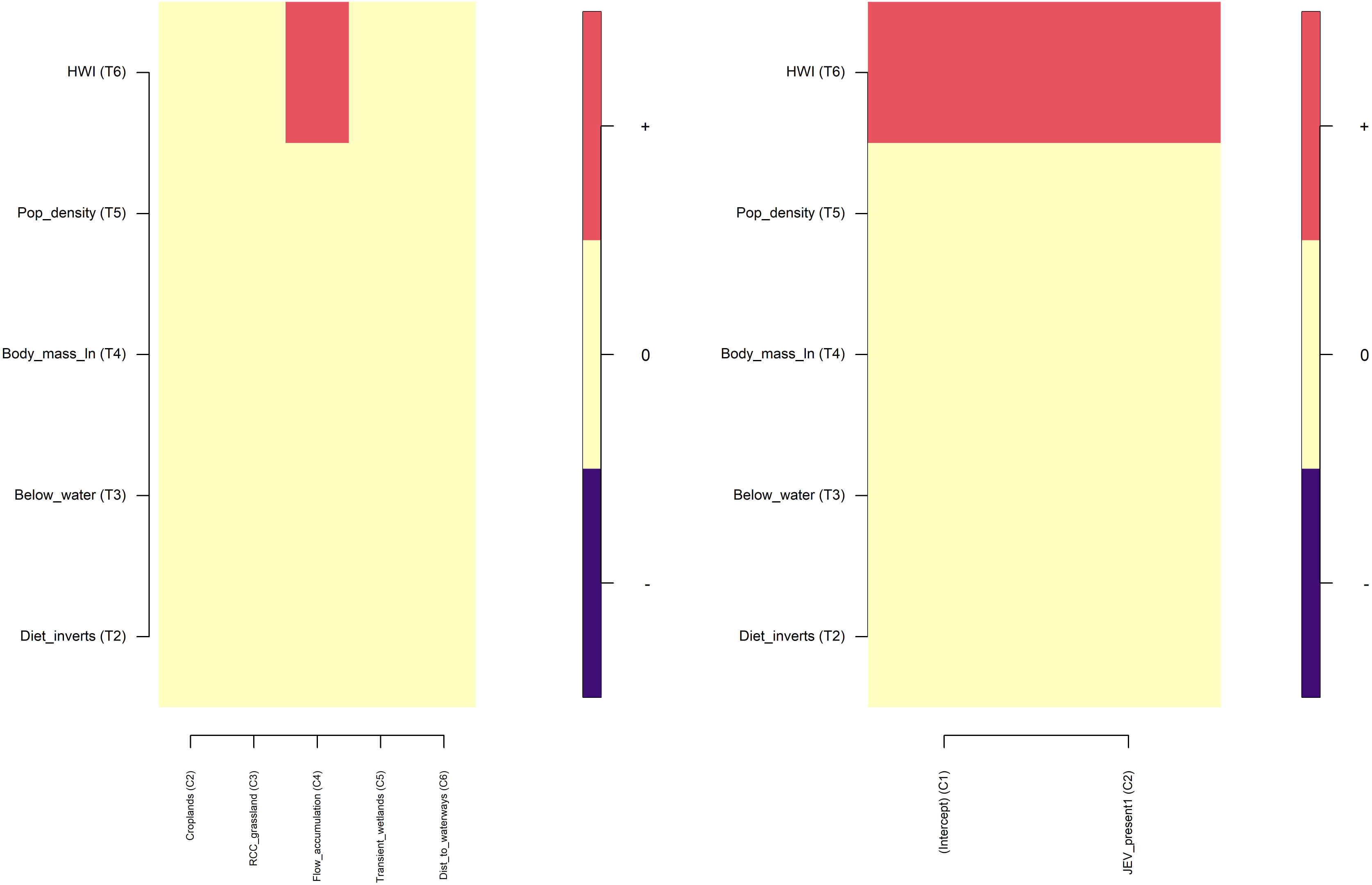
The association between species traits and the landscape features, representing modulation of species “responses” to the landscape features, under two model structures using hierarchical modelling of species communities (HMSC). The first HMSC model (left panel) shows associations as positive, negative, or neutral “modulators” for each trait on the y-axis of each landscape feature associated with JEV outbreaks on the x-axis. The second HMSC model (right panel) shows these associations for JEV outbreak occurrence only and no other landscape features.

Interestingly, further insight about the broad associations between hand-wing index and species presence in JEV-associated landscapes in Australia and across the CIPBR was provided when this trait was mapped to the waterbird phylogeny (Figure 6), and both Blomberg’s K (K = 0.345; p = 0.001) and Pagel’s λ (λ = 0.874; p < 0.00001) showed that hand-wing index was phylogenetically correlated across waterbird species. Although the ardeid family showed a moderate to high hand-wing index, the anatid, threskiornothid, and phalacrocoracid families showed substantively higher hand-wing index, as did a couple of individual stork species (Ciconiidae) and pelicans (Pelecanidae).

**Figure 6.**
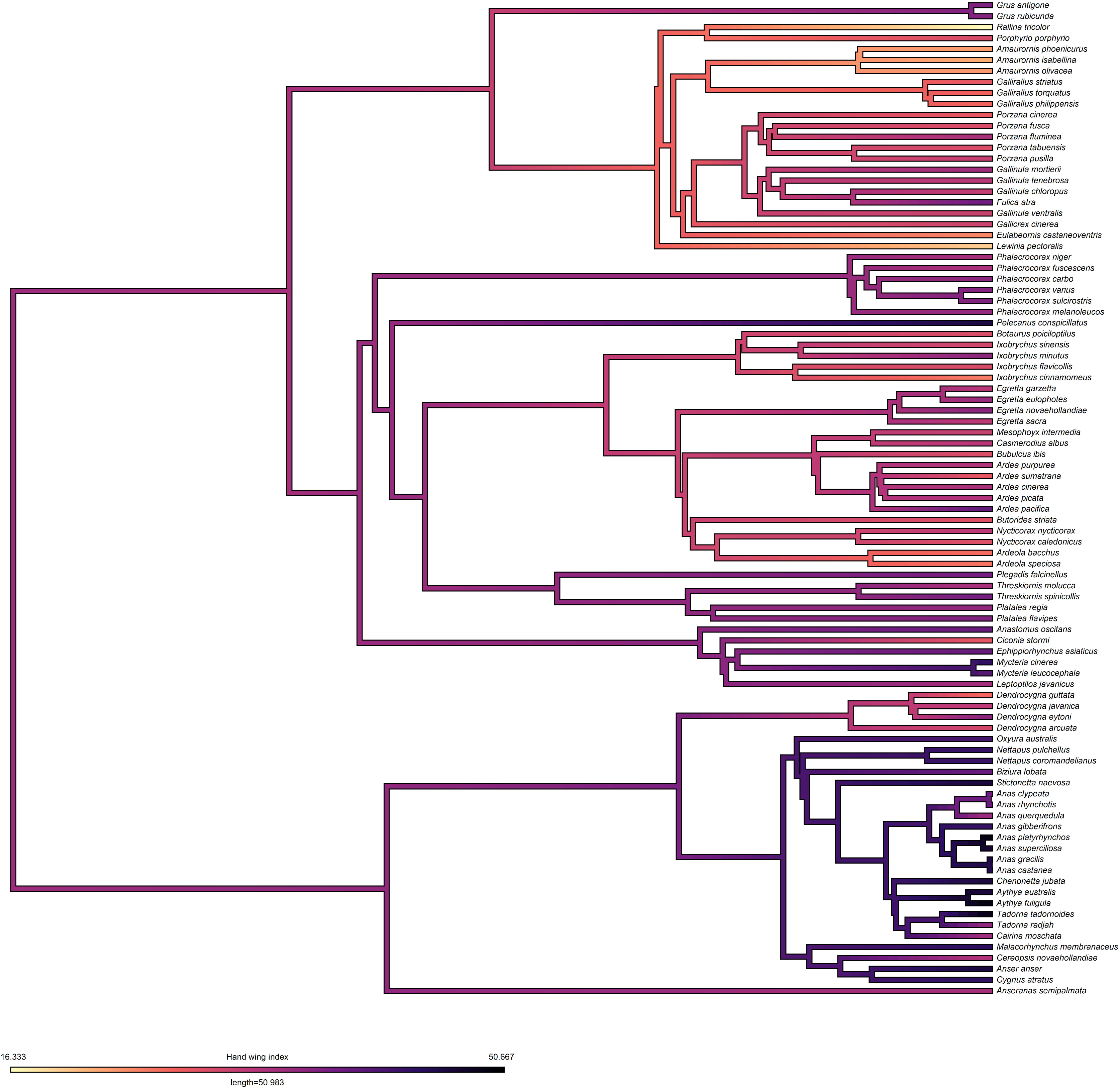
Mapping of hand-wing index onto the phylogenetic tree for 91 waterbird species evaluated across the Central Indo-Pacific biogeographical region.

## Discussion

This study describes waterbird presence and the functional structure of species pools at landscape scale to inform JEV surveillance in high-risk landscapes for JEV circulation in Australia, as well as providing evidence that these associations might reflect the functional biogeography (hand- wing index) of species presence relative to JEV circulation.

Although several ardeid species were strongly associated with the distribution of JEV detections in Australia and across the CIPBR, species presence was neither limited to nor dominated by ardeids, challenging conventional wisdom and suggesting that JEV surveillance should include a broader range of potential waterbird hosts. Hand-wing index was associated with species presence in landscapes demarcated by JEV occurrence in both Australia and the broader CIPBR, and simultaneously demonstrated an association with species presence in areas with transient wetlands in Australia, which may indicate functional modulation of species with respect to this high-risk JEV- associated landscape feature. Contrary to expectation, increasing functional divergence among the species pools was associated with JEV detections. For the purposes of wildlife surveillance development in Australia, these findings suggest that whilst high-risk JEV landscapes can vary with respect to the waterbird species present, JEV surveillance across broad regions of eastern and southern Australia can be informed using a combination of landscape structure and targeting of individual species in these landscapes (see Table S2 for full list of associated species). Moreover, a more nuanced consideration of the functional biogeography of waterbird species across Australia and the CIPBR suggests that broader, regional surveillance development may benefit from considering target species with trait profiles that favour dispersal.

The extraordinarily broad geographic distribution of JEV outbreaks in Australia makes developing a representative, systematic sampling strategy to answer key maintenance host ecology questions exceedingly challenging. This study informs the development of such a sampling strategy, describing waterbird presence as a function of the available species pool and enabling species presence to be directly evaluated with respect to the landscape structure associated with JEV outbreaks, and therefore that landscape structure can inform species selection for surveillance. Moreover, by further considering species presence across the regional as whole, this study was able to interrogate functional traits relevant to pathogen transmission within the context of known regional transmission of JEV. The current study identified several species that were individually associated with JEV in Australia and across the CIPBR and found that many of these species represent families other than the Ardeidae, suggesting that conventional understanding JEV maintenance hosts may be too narrow in scope. Moreover, in the Australian context, species presence is likely to be heterogeneous across the different landscape features associated with JEV outbreaks, and these associations can reflect differences in functional traits (e.g. dispersal capacity) that are relevant to both arbovirus transmission and response to environmental variation at landscape scale (i.e. flooding).

Associations between the dominant traits of species present in the landscape species pool may provide insight into potential drivers of species structure in landscapes delineating JEV risk. For example, landscapes associated with JEV occurrence were also characterised by species demonstrating a high level of dispersal capacity, as indicated by greater hand-wing index. This may indicate the importance of dispersal among species in the context of seasonal or transient wetlands, which were strongly associated with species presence in the current study, and Australian JEV outbreaks previously(Walsh et al., 2023). Such a scenario is supported by analyses of population structure among birds in Australia following La Niña precipitation anomalies(Englert Duursma et al., 2018; Purnell, 2022), and reflects more general evidence for the importance of hand-wing index as an indicator of dispersal capacity in response to climate gradients and habitat fragmentation(Kennedy et al., 2016; Sheard et al., 2020; Weeks et al., 2023). Moreover, dispersal capacity as indicated by hand-wing index has also been shown to be associated with bird host status and landscape-level outbreak occurrence, albeit for a different viral pathogen but one that, nevertheless, involves waterbirds in agricultural-wetland mosaics (Yin et al., 2023). Furthermore, the strong association between hand-wing index and JEV landscapes in the current study extended to the whole of the CIPBR, although this investigation was not able to deconstruct the composition and configuration of these landscapes for the region as a whole as it did for Australia, specifically. Interestingly, the two broad-scale multiple PPM approaches (Australia and CIPBR) generally agreed with respect to the best fitting models comprising hand-wing index and MPD, and both also univariately identified associations with population density and foraging strategy. The association between JEV detection and higher population density is not surprising as this may reflect pathogen transmission dynamics(Adler et al., 2008; Begon et al., 2002; Thrall et al., 1995), but it did not persist after having accounted for hand-wing index, MPD and foraging strategy in the Australian context and hand-wing index and MPD in the CIPBR context. It should also be noted that the association between hand-wing index and species presence in high-risk landscapes was not an artifact of body mass because these traits were not correlated (Pearson’s r = -0.21), which is as expected since hand-wing index accounts for body size(Lockwood et al., 1998). While these findings are intriguing, it is critical to emphasise that they are only correlative of possible drivers of community assembly at landscape scale and do not describe realised community structure or specific modulation of environmental (or biotic) filtering by species traits. It is not possible to infer local community composition or functional responses to environmental conditions from the data presented here. Rather, the true utility of this work is to describe preliminary, evidence-based, configurations of waterbird species pools for the coupling of high-risk JEV landscape structure with potential maintenance hosts across extraordinarily broad geographic extent, and thereby inform the development of broad-scale sampling strategies for One Health JEV surveillance, specifically for Australia but with potential impact for the broader region.

Beyond the implications of these findings for potential wild bird surveillance programs, there may be potential utility for the design of arbovirus surveillance programs by local health authorities in Australia. Monitoring sites is often informed by historic arbovirus activity or according to potential mosquito and waterbird habitats. Consideration of wildlife hosts, and concomitant traits, in specific landscapes may provide additional information in identifying appropriate arbovirus surveillance locations. Nevertheless, a thorough investigation of mosquito communities and vector-host interactions between mosquito and waterbird communities, particularly across heterogenous features of high-risk landscapes, will also be essential for maximally beneficial surveillance output. The current study was not able to interrogate vector communities specifically, and as such this remains a key knowledge gap in the infection ecology of JEV.

In addition to the specific constraints to interpretation of the results described above, further description of additional limitations is included below. Firstly, both the recording of bird observations and the reporting of piggery outbreaks in Australia and other JEV detections across the CIPRB may be affected by reporting biases. We have corrected for potential biases, respectively, by using spatial block partitioning in the construct of the SDMs, and sampling background points for JEV occurrence proportional to mean pig density, as described above. Nevertheless, it is possible that residual bias may persist. Moreover, it is important to note that the sample of JEV detections across the CIPBR is small and, despite this sample being both reflective of piggeries and corrected for potential reporting bias, likely reflects quite different surveillance systems to those that generated the reporting for piggery outbreaks in Australia. Therefore, we emphasise that this sample is in no way intended to be representative of JEV presence across the region. Rather, it is intended as a rudimentary comparison to the mapping of waterbird species in Australia to evaluate whether there may be a potential shared biogeography of waterbird hosts across the broader region. The primary practical application of the findings for wildlife monitoring in JEV surveillance are nevertheless limited to Australia. Secondly, notwithstanding the inclusion of each of the waterbird species found in the CIPBR in the habitat suitability modelling, it is important to note that suitability can be interpreted with respect to the fundamental niche only, because it does not include direct observational evaluation of biotic interactions to more formally define the realised niche or subsequent community structure. We must acknowledge that neither local biotic interactions nor dispersal history are represented in this framework and should be considered in future work. Thirdly, the climate measures used in the waterbird habitat suitability models were derived from decadal averages from 1970 to 2000, which assumes homogeneity over this period as well as over the period of bird observations from 2010 to 2020. Although the current study used these features to model the influence of climate features on habitat suitability rather than specific weather events, which is appropriate, we do note that, due to the rapid increase in global temperatures over the last 20 years, climatic means could have already shifted by 2020 from the baseline represented by the period 1970-2000. Finally, previous work showed that the piggery outbreaks were not associated with feral pigs(Walsh et al., 2023). Nevertheless, JEV has been identified in feral pigs in Australia, albeit only very sporadically(Australian Veterinary Association, 2023). As such, it is possible that there may be multiple landscapes of JEV risk, both within Australia and the broader CIPBR: those pertaining to domestic piggeries and those pertaining to feral pig habitat. If the latter are in fact present and represent impactful nidi of JEV circulation, then different surveillance programs targeting different maintenance host communities may be required.

## Conclusions

This study has identified a breadth of waterbird species in association with particular configurations of landscape risk that extends well beyond the Ardeidae family, the taxon to which epidemiological and ecological investigations of JEV maintenance hosts are typically constrained. Moreover, the findings relate species presence in JEV-associated landscapes to waterbird functional biogeography across the CIPBR, highlighting dispersal capacity in particular. Although preliminary, it is anticipated that together these results can help to inform an evidence-based approach to JEV surveillance development that can provide a more efficient and productive yield than a simple random or convenience-based approach to maintenance host sampling, particularly given the wide geographic extent in which JEV outbreaks occurred across the region. As wild bird monitoring progresses and local community structure becomes better articulated with respect to JEV circulation, we expect that the scope of JEV maintenance understanding will be updated to reflect currently unrecognised local heterogeneities in landscape and community composition, as well as novel maintenance hosts.

## Acknowledgements

We thank Australian Pork Limited for supporting (non-financially) this research. We thank SunPork Group for information about case farms. We thank Richard Bradhurst from the University of Melbourne who facilitated the sharing of the data products.

## Data Availability Statement

All data described in this study are publicly available for download at the cited sources in the text except the pig density data, which contains sensitive information to the Australian agricultural industry (e.g. locations of piggeries). Provision of this data for third parties requires their application to Australian Pork Limited.

## Author statement

Conceptualization: Michael Walsh, Cameron Webb, Victoria Brookes; Data curation: Michael Walsh and Victoria Brookes; Formal analysis: Michael Walsh; Investigation: Michael Walsh, Cameron Webb, Victoria Brookes; Methodology: Michael Walsh, Victoria Brookes; Validation: Michael Walsh, Cameron Webb, Victoria Brookes; Visualization: Michael Walsh; Writing - original draft: Michael Walsh, Writing - review & editing: Michael Walsh, Cameron Webb, Victoria Brookes

