## Supplemental material for "Landscapes associated with Japanese encephalitis virus reflect the functional biogeography of waterbird species across Australia and the Central Indo-Pacific region"

Table S1. Waterbird species comparisons based on ensemble species distribution models. Each species listed is presented with their associated number of observations in the field (and the number of observations used for analysis after thinning in parentheses), their model omission rate (OR), true skill statistic (TSS), and area under the receiver operating characteristic curve (AUC).

| Bird species grouped by Family | Number of observations | OR | TSS | AUC |
|---|---|---|---|---|
| **Ardeidae** | | | | |
| *Casmerodius albus (Ardea alba; Ardea modesta)** | 172534 (20156) | 0.126 | 0.753 | 0.944 |
| *Ardea cinerea* | 6108 (1286) | 0.115 | 0.758 | 0.939 |
| *Ardea pacifica* | 62619 (17236) | 0.144 | 0.692 | 0.916 |
| *Ardea purpurea* | 11360 (2197) | 0.154 | 0.670 | 0.911 |
| *Ardea sumatrana* | 2851 (859) | 0.114 | 0.731 | 0.926 |
| *Ardeola bacchus* | 1211 (358) | 0.096 | 0.723 | 0.921 |
| *Ardeola speciosa* | 6379 (1458) | 0.124 | 0.669 | 0.908 |
| *Botaurus poiciloptilus* | 2158 (518) | 0.0538 | 0.809 | 0.950 |
| *Bubulcus ibis** | 92098 (14689) | 0.0969 | 0.792 | 0.959 |
| *Butorides striata* | 35407 (5726) | 0.0891 | 0.820 | 0.968 |
| *Ixobrychus flavicollis* (*Dupetor flavicollis*) | 3968 (1222) | 0.097 | 0.734 | 0.933 |
| *Egretta eulophotes* | 1037 (241) | 0.090 | 0.748 | 0.937 |
| *Egretta garzetta* | 85177 (11159) | 0.113 | 0.765 | 0.951 |
| *Mesophoyx intermedia* (*Egretta intermedia; Ardea intermedia*) | 72270 (9288) | 0.124 | 0.747 | 0.941 |
| *Egretta novaehollandiae* | 267840 (35103) | 0.106 | 0.771 | 0.948 |
| *Egretta picata* | 7136 (947) | 0.0678 | 0.848 | 0.971 |
| *Egretta sacra* | 22038 (4444) | 0.090 | 0.819 | 0.966 |
| *Ixobrychus cinnamomeus* | 3539 (912) | 0.200 | 0.636 | 0.888 |
| *Ixobrychus sinensis* | 4875 (1087) | 0.135 | 0.749 | 0.933 |
| *Ixobrychus minutus (Ixobrychus dubius)* | 1832 (314) | 0.067 | 0.856 | 0.968 |
| *Nycticorax caledonicus* | 36488 (6953) | 0.142 | 0.730 | 0.937 |
| *Nycticorax nycticorax* | 8004 (1289) | 0.106 | 0.737 | 0.925 |
| **Anatidae** | | | | |
| *Anas castanea* | 147348 (10964) | 0.0409 | 0.875 | 0.979 |
| *Anas gibberifrons* | 1560 (349) | 0.140 | 0.684 | 0.886 |
| *Anas gracilis* | 217162 (21620) | 0.158 | 0.715 | 0.926 |
| *Anas platyrhynchos* | 24421 (2758) | 0.0583 | 0.866 | 0.977 |
| *Anas superciliosa* | 417907 (37537) | 0.100 | 0.790 | 0.954 |
| *Anastomus oscitans* | 1608 (338) | 0.086 | 0.836 | 0.965 |
| *Anser anser* | 3554 (540) | 0.0469 | 0.880 | 0.974 |
| *Anseranas semipalmata* | 66065 (6709) | 0.0762 | 0.855 | 0.974 |
| *Aythya australis* | 129262 (11639) | 0.139 | 0.747 | 0.938 |
| *Aythya fuligula* | 662 (103) | 0.0780 | 0.785 | 0.931 |
| *Biziura lobata* | 31349 (3371) | 0.0841 | 0.831 | 0.966 |
| *Cairina moschata* | 6601 (1054) | 0.0797 | 0.840 | 0.967 |
| *Cereopsis novaehollandiae* | 14142 (1495) | 0.0241 | 0.862 | 0.975 |
| *Chenonetta jubata* | 285930 (36218) | 0.0848 | 0.795 | 0.954 |
| *Cygnus atratus* | 186734 (15800) | 0.0913 | 0.818 | 0.966 |
| *Dendrocygna arcuate* | 21592 (2477) | 0.115 | 0.764 | 0.943 |
| *Dendrocygna eytoni* | 27769 (4495) | 0.173 | 0.688 | 0.921 |
| *Dendrocygna guttata* | 1242 (246) | 0.0874 | 0.704 | 0.895 |
| *Dendrocygna javanica* | 1198 (314) | 0.160 | 0.679 | 0.882 |
| *Malacorhynchus membranaceus* | 51148 (5498) | 0.197 | 0.652 | 0.904 |
| *Nettapus coromandelianus* | 5037 (757) | 0.127 | 0.785 | 0.950 |
| *Nettapus pulchellus* | 12567 (1108) | 0.122 | 0.773 | 0.951 |
| *Oxyura australis* | 21350 (1659) | 0.0606 | 0.819 | 0.959 |
| *Radjah radjah (Tadorna radjah)* | 14997 (1773) | 0.0663 | 0.843 | 0.972 |
| *Spatula clypeata* | 815 (84) | 0.0546 | 0.883 | 0.955 |

| Bird species | Number of observations | OR | TSS | AUC |
|---|---|---|---|---|
| *Spatula querquedula* | 577 (81) | 0.141 | 0.784 | 0.920 |
| *Spatula rhynchotis* | 45225 (4360) | 0.0662 | 0.817 | 0.964 |
| *Stictonetta naevosa* | 19094 (1785) | 0.125 | 0.756 | 0.942 |
| *Tadorna tadornoides* | 57949 (9395) | 0.0654 | 0.837 | 0.966 |
| **Rallidae** | | | | |
| *Amaurornis isabellina* | 507 (100) | 0.104 | 0.697 | 0.860 |
| *Amaurornis olivacea* | 5077 (1120) | 0.101 | 0.762 | 0.940 |
| *Amaurornis phoenicurus* | 14029 (2832) | 0.188 | 0.572 | 0.857 |
| *Fulica atra* | 236742 (15678) | 0.105 | 0.798 | 0.957 |
| *Gallicrex cinerea* | 1087 (264) | 0.133 | 0.759 | 0.928 |
| *Gallinula chloropus* | 4571 (867) | 0.159 | 0.681 | 0.893 |
| *Gallinula mortierii* | 14929 (1551) | 0.0840 | 0.785 | 0.946 |
| *Gallinula tenebrosa* | 209156 (13326) | 0.0604 | 0.850 | 0.974 |
| *Gallinula ventralis* | 21397 (4214) | 0.173 | 0.656 | 0.908 |
| *Gallirallus castaneoventris* | 510 (101) | 0.000 | 0.954 | 0.968 |
| *Gallirallus philippensis* | 28827 (5134) | 0.113 | 0.794 | 0.959 |
| *Gallirallus striatus* | 481 (197) | 0.240 | 0.589 | 0.837 |
| *Gallirallus torquatus* | 4529 (924) | 0.109 | 0.657 | 0.878 |
| *Lewinia pectoralis* | 4660 (968) | 0.0643 | 0.835 | 0.963 |
| *Porphyrio indicus* | 551 (152) | 0.102 | 0.647 | 0.882 |
| *Porphyrio melanotus* | 191072 (11214) | 0.0847 | 0.837 | 0.969 |
| *Porzana cinerea* | 6273 (959) | 0.112 | 0.727 | 0.934 |
| *Porzana fluminea* | 13004 (1660) | 0.103 | 0.774 | 0.953 |
| *Porzana fusca* | 520 (197) | 0.128 | 0.601 | 0.872 |
| *Porzana pusilla* | 6920 (1016) | 0.0839 | 0.786 | 0.950 |
| *Porzana tabuensis* | 10930 (1510) | 0.0921 | 0.829 | 0.966 |
| *Rallina tricolor* | 1378 (230) | 0.191 | 0.703 | 0.908 |
| **Phalacrocoracidae** | | | | |
| *Microcarbo melanoleucos* | 270440 (26275) | 0.113 | 0.781 | 0.953 |
| *Microcarbo niger* | 699 (152) | 0.079 | 0.820 | 0.945 |
| *Phalacrocorax carbo* | 113515 (14517) | 0.0800 | 0.831 | 0.971 |
| *Phalacrocorax fuscescens* | 12871 (2075) | 0.0396 | 0.906 | 0.983 |
| *Phalacrocorax sulcirostris* | 208014 (20119) | 0.109 | 0.787 | 0.957 |
| *Phalacrocorax varius* | 83644 (11821) | 0.0916 | 0.819 | 0.968 |
| **Threskiornithidae** | | | | |
| *Platalea flavipes* | 47110 (8048) | 0.105 | 0.769 | 0.947 |
| *Platalea regia* | 92577 (9742) | 0.113 | 0.787 | 0.955 |
| *Plegadis falcinellus* | 30491 (3983) | 0.126 | 0.739 | 0.935 |
| *Threskiornis Molucca* | 300828 (24823) | 0.0747 | 0.829 | 0.969 |
| *Threskiornis spinicollis* | 157646 (24537) | 0.109 | 0.768 | 0.946 |
| **Gruidae** | | | | |
| *Grus Antigone* | 2263 (561) | 0.106 | 0.734 | 0.921 |
| *Grus rubicunda* | 46877 (7789) | 0.148 | 0.677 | 0.910 |
| **Cinconiidae** | | | | |
| *Ciconia stormi* | 1727 (252) | 0.101 | 0.667 | 0.879 |
| *Ephippiorhynchus asiaticus* | 17486 (4275) | 0.090 | 0.815 | 0.964 |
| *Leptoptilos javanicus* | 2289 (513) | 0.179 | 0.619 | 0.868 |
| *Mycteria cinerea* | 526 (156) | 0.142 | 0.662 | 0.861 |
| *Mycteria leucocephala* | 814 (326) | 0.093 | 0.786 | 0.935 |
| **Pelecanidae** | | | | |
| *Pelecanus conspicillatus* | 217875 (21721) | 0.103 | 0.805 | 0.963 |

* *Ardea modesta* was previously considered a subspecies of *Casmerodius albus (Ardea alba)* and *Bubulcus coromandus* was previously considered a subspecies of *Bubulcus ibis*. These were ultimately combined to estimate *Casmerodius albus* and *Bubulcus coromandus* habitat suitabilities, respectively. Blue = endemic or introduced to Australia; orange = occurs across CIPBR, excluding Australia; black = occurs across CIPBR, including Australia.

Table S2. Univariable regression coefficients and 95% confidence intervals for those species demonstrating associations between their habitat suitability Japanese encephalitis virus outbreaks in Australia in 2021-2022. Coefficients are derived from inhomogeneous Poisson models.

| **Landscape species pool abundance** | **AIC** | **Estimate** | **95% confidence interval** |
|---|---|---|---|
| Null model | 243.05 | | |
| *Casmerodius albus* (*Ardea alba*) | 221.52 | 0.017 | 0.010 – 0.023 |
| *Ardea pacifica* | 207.33 | 0.019 | 0.013 – 0.026 |
| *Botaurus poiciloptilus* | 205.98 | 0.026 | 0.019 – 0.033 |
| *Mesophoyx intermedia (Egretta intermedia)* | 240.02 | 0.010 | 0.001 – 0.016 |
| *Egretta novaehollandiae* | 214.12 | 0.018 | 0.011 – 0.024 |
| *Ixobrychus minutus* (*Ixobrychus dubius*) | 206.50 | 0.025 | 0.018 – 0.032 |
| *Nycticorax caledonicus* | 221.91 | 0.017 | 0.010 – 0.023 |
| *Anas castanea* | 228.32 | 0.017 | 0.009 – 0.024 |
| *Anas gracilis* | 185.34 | 0.025 | 0.018 – 0.032 |
| *Anas platyrhynchos* | 237.78 | 0.012 | 0.004 – 0.021 |
| *Anas superciliosa* | 217.33 | 0.017 | 0.010 – 0.023 |
| *Aythya australis* | 216.74 | 0.017 | 0.011 – 0.023 |
| *Biziura lobata* | 220.10 | 0.021 | 0.014 – 0.029 |
| *Cereopsis novaehollandiae* | 231.34 | 0.025 | 0.014 – 0.035 |
| *Chenonetta jubata* | 219.33 | 0.016 | 0.010 – 0.022 |
| *Cygnus atratus* | 223.62 | 0.017 | 0.010 – 0.024 |
| *Dendrocygna eytoni* | 211.19 | 0.018 | 0.012 – 0.025 |
| *Malacorhynchus membranaceus* | 192.71 | 0.024 | 0.017 – 0.031 |
| *Oxyura australis* | 210.53 | 0.022 | 0.016 – 0.029 |
| *Spatula clypeata* | 219.24 | 0.053 | 0.037 – 0.069 |
| *Spatula rhynchotis* | 209.33 | 0.022 | 0.016 – 0.029 |
| *Stictonetta naevosa* | 207.22 | 0.021 | 0.015 – 0.028 |
| *Tadorna tadornoides* | 193.46 | 0.023 | 0.017 – 0.029 |
| *Fulica atra* | 220.85 | 0.016 | 0.010 – 0.023 |
| *Gallinula tenebrosa* | 240.08 | 0.009 | 0.001 – 0.016 |
| *Gallinula ventralis* | 159.22 | 0.030 | 0.023 – 0.036 |
| *Porphyrio melanotus* | 239.54 | 0.009 | 0.001 – 0.016 |
| *Porzana fluminea* | 181.74 | 0.028 | 0.022 – 0.035 |
| *Porzana pusilla* | 200.63 | 0.024 | 0.017 – 0.030 |
| *Porzana tabuensis* | 226.19 | 0.017 | 0.010 – 0.024 |
| *Microcarbo melanoleucos* | 226.87 | 0.014 | 0.008 – 0.021 |
| *Phalacrocorax carbo* | 212.65 | 0.021 | 0.014 – 0.028 |
| *Phalacrocorax fuscescens* | 234.55 | 0.026 | 0.013 – 0.040 |
| *Phalacrocorax sulcirostris* | 227.87 | 0.015 | 0.008 – 0.021 |
| *Phalacrocorax varius* | 234.98 | 0.014 | 0.006 – 0.022 |
| *Platalea flavipes* | 191.22 | 0.024 | 0.018 – 0.031 |
| *Platalea regia* | 225.09 | 0.017 | 0.010 – 0.023 |
| *Plegadis falcinellus* | 214.83 | 0.019 | 0.013 – 0.026 |
| *Threskiornis Molucca* | 224.52 | 0.016 | 0.009 – 0.022 |
| *Threskiornis spinicollis* | 216.51 | 0.017 | 0.011 – 0.023 |
| *Grus rubicunda* | 229.97 | 0.014 | 0.007 – 0.021 |
| *Pelecanus conspicillatus* | 218.74 | 0.019 | 0.012 – 0.025 |

Table S3. Univariable regression coefficients and 95% confidence intervals for those species demonstrating associations between their habitat suitability and Japanese encephalitis virus detections across the broader CIPBR (including reported outbreaks in Australia in 2021-2022). Coefficients are derived from inhomogeneous Poisson models. Species highlighted in red indicate species associated with JEV detections across the whole CIPBR but not with the Australian outbreaks alone (Table S2).

| **Landscape species pool abundance** | **AIC** | **Estimate** | **95% confidence interval** |
|---|---|---|---|
| Null model | 838.56 | | |
| *Casmerodius albus* (*Ardea alba*) | 475.80 | 0.026 | 0.020 – 0.032 |
| *Ardea pacifica* | 445.43 | 0.025 | 0.020 – 0.030 |
| *Botaurus poiciloptilus* | 469.81 | 0.032 | 0.026 – 0.039 |
| *Mesophoyx intermedia (Egretta intermedia)* | 511.06 | 0.019 | 0.012 – 0.026 |
| *Egretta garzetta* | 520.41 | 0.015 | 0.007 – 0 .024 |
| *Egretta novaehollandiae* | 455.08 | 0.026 | 0.021 – 0.031 |
| *Ixobrychus minutus* (*Ixobrychus dubius*) | 462.16 | 0.033 | 0.027 – 0.039 |
| *Nycticorax caledonicus* | 471.48 | 0.026 | 0.020 – 0.031 |
| *Anas castanea* | 491.58 | 0.026 | 0.020 – 0.033 |
| *Anas gracilis* | 423.66 | 0.029 | 0.024 – 0.034 |
| *Anas platyrhynchos* | 504.08 | 0.026 | 0.018 – 0.034 |
| *Anas superciliosa* | 460.95 | 0.025 | 0.020 – 0.030 |
| *Anser anser* | 517.20 | 0.020 | 0.011 – 0.028 |
| *Aythya australis* | 462.70 | 0.025 | 0.020 – 0.030 |
| *Biziura lobata* | 488.13 | 0.028 | 0.021 – 0.034 |
| *Cairina moschata* | 517.91 | 0.019 | 0.010 – 0.028 |
| *Cereopsis novaehollandiae* | 507.85 | 0.034 | 0.024 – 0.045 |
| *Chenonetta jubata* | 460.61 | 0.025 | 0.020 – 0.030 |
| *Cygnus atratus* | 482.17 | 0.025 | 0.019 – 0.031 |
| *Dendrocygna eytoni* | 454.56 | 0.026 | 0.021 – 0.031 |
| *Malacorhynchus membranaceus* | 437.70 | 0.028 | 0.023 – 0.033 |
| *Oxyura australis* | 470.26 | 0.029 | 0.023 – 0.035 |
| *Spatula clypeata* | 522.00 | 0.020 | 0.008 – 0.032 |
| *Spatula rhynchotis* | 465.70 | 0.029 | 0.023 – 0.035 |
| *Stictonetta naevosa* | 461.45 | 0.027 | 0.022 – 0.033 |
| *Tadorna tadornoides* | 448.76 | 0.029 | 0.024 – 0.034 |
| *Fulica atra* | 471.45 | 0.025 | 0.020 – 0.031 |
| *Gallinula chloropus* | 473.56 | 0.027 | 0.021 – 0.033 |
| *Gallinula tenebrosa* | 504.69 | 0.022 | 0.015 – 0.029 |
| *Gallinula ventralis* | 405.75 | 0.031 | 0.027 – 0.036 |
| *Gallirallus philippensis* | 511.38 | 0.017 | 0.011 – 0.024 |
| *Gallirallus torquatus* | 518.54 | 0.020 | 0.010 – 0.029 |
| *Porphyrio melanotus* | 505.95 | 0.020 | 0.014 – 0.027 |
| *Porzana fluminea* | 425.69 | 0.035 | 0.029 – 0.040 |
| *Porzana pusilla* | 444.14 | 0.032 | 0.026 – 0.037 |
| *Porzana tabuensis* | 485.92 | 0.026 | 0.020 – 0.032 |
| *Microcarbo melanoleucos* | 475.66 | 0.025 | 0.019 – 0.030 |
| *Phalacrocorax carbo* | 464.15 | 0.030 | 0.024 – 0.035 |
| *Phalacrocorax fuscescens* | 515.52 | 0.032 | 0.020 – 0.045 |
| *Phalacrocorax sulcirostris* | 483.07 | 0.025 | 0.019 – 0.031 |
| *Phalacrocorax varius* | 499.56 | 0.026 | 0.019 – 0.034 |
| *Platalea flavipes* | 431.94 | 0.030 | 0.025 – 0.035 |
| *Platalea regia* | 477.62 | 0.027 | 0.021 – 0.033 |
| *Plegadis falcinellus* | 463.55 | 0.028 | 0.022 – 0.033 |
| *Threskiornis Molucca* | 473.81 | 0.026 | 0.021 – 0.032 |
| *Threskiornis spinicollis* | 456.99 | 0.026 | 0.020 – 0.031 |
| *Grus rubicunda* | 495.52 | 0.021 | 0.016 – 0.027 |
| *Mycteria leucocephala* | 526.55 | 0.029 | 0.007 – 0.051 |
| *Pelecanus conspicillatus* | 468.32 | 0.029 | 0.023 – 0.035 |

Figure S1. Landscape features associated with Japanese encephalitis virus (JEV) piggery outbreaks (Walsh, M.G., Webb, C., Brookes, V. One Health, Volume 16, June 2023.). Overlaid points show the locations of the JEV outbreaks.

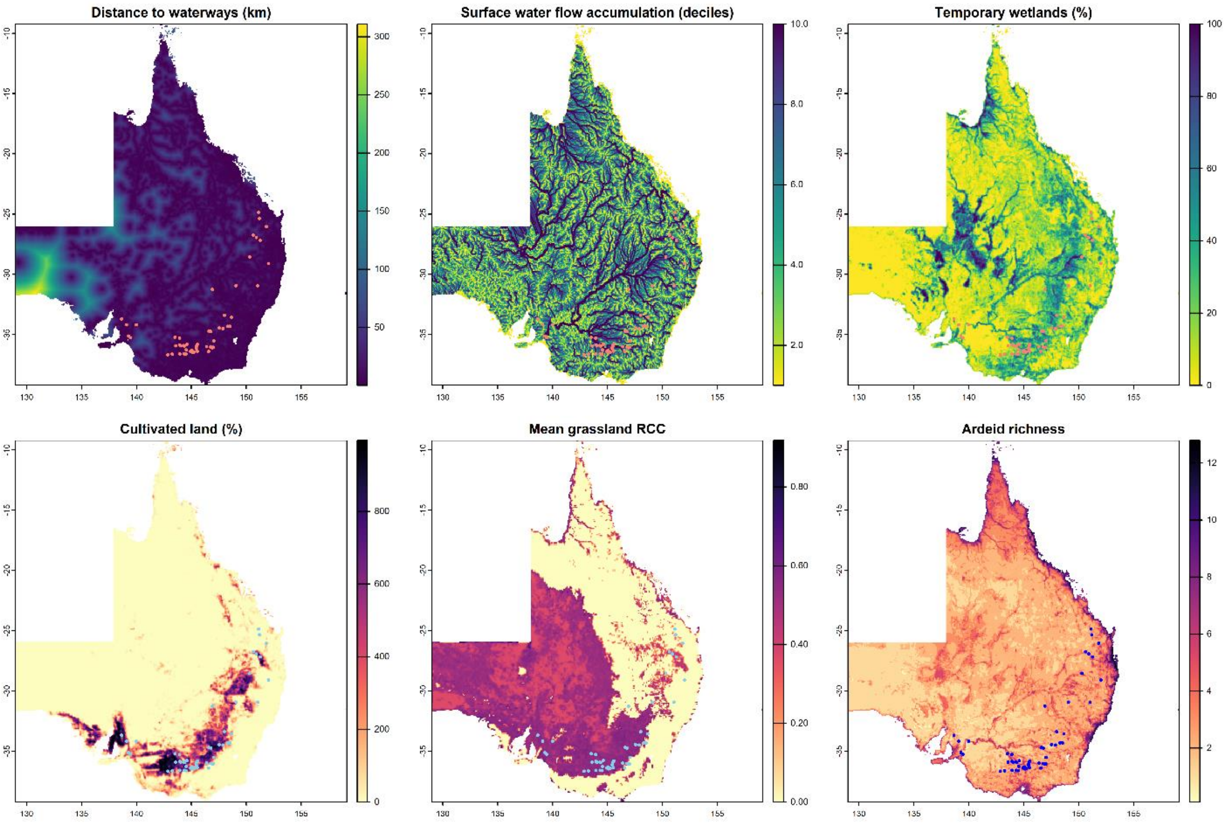


RCC: related circumscribing circle

Table S4. Univariable regression coefficients and 95% confidence intervals for the associations between Japanese encephalitis virus outbreaks and each species trait as derived from simple univariate inhomogeneous Poisson models for Australia alone (A) and the Central Indo-Pacific biogeographical region (CIPBR) as a whole (B).

| **Landscape-scale species pool-weighted mean traits** | **AIC** | **Estimate** | **95% confidence interval** |
|---|---|---|---|
| **A. Australia-specific univariate models** | | | |
| *Null model* | 236.75 | | |
| Percentage of diet – fish | 233.83 | 0.001 | -0.033 – 0.034 |
| Percentage of diet – invertebrates | 233.81 | -0.002 | -0.031 – 0.026 |
| Percentage of diet – plants | 233.58 | 0.007 | -0.021 – 0.035 |
| Foraging strategy – underwater | 228.08 | 0.039 | 0.006 – 0.073 |
| Foraging strategy – water surface | 233.83 | 0.001 | -0.021 – 0.022 |
| Foraging strategy – ground | 230.05 | -0.024 | -0.049 – 0.001 |
| Body mass (ln(g)) | 232.62 | -0.375 | -1.083 – 0.332 |
| Egg mass (g) | 232.75 | -0.007 | -0.021 – 0.007 |
| Hand-wing index | 220.13 | 0.190 | 0.095 – 0.285 |
| Population density (birds/km$^2$) | 220.33 | 0.345 | 0.177 – 0.514 |
| Mean pairwise dissimilarity | 227.69 | 6.564 | 0.490 – 12.637 |
| **B. CIPBR univariate models** | | | |
| *Null model* | 838.56 | | |
| Percentage of diet – fish | 551.62 | -0.024 | -0.043 – -0.006 |
| Percentage of diet – invertebrates | 549.05 | -0.040 | -0.064 – -0.016 |
| Percentage of diet – plants | 525.95 | 0.042 | 0.030 – 0.054 |
| Foraging strategy – underwater | 552.52 | 0.028 | 0.006 – 0.050 |
| Foraging strategy – water surface | 557.41 | -0.008 | -0.025 – 0.008 |
| Foraging strategy – ground | 557.60 | -0.008 | -0.026 – 0.010 |
| Body mass (ln(g)) | 556.81 | 0.283 | -0.150 – 0.715 |
| Egg mass (g) | 554.69 | 0.012 | 0.001 – 0.022 |
| Hand-wing index | 496.35 | 0.196 | 0.150 – 0.243 |
| Population density (birds/km$^2$) | 535.15 | 0.185 | 0.311 – 1.290 |
| Mean pairwise dissimilarity | 497.10 | 17.868 | 12.738 – 22.999 |

Red = statistically significant univariate associations.
